# Giant bacteria in gut microbiomes

**DOI:** 10.64898/2026.09.30.750856

**Authors:** Yiang Lyu, Marit Grüner, Zhen Han, Aron Klein, Luca Zoccarato, Alon Daya, Hyun-Dong Chang, Mina Bizic, Danny Ionescu

## Abstract

Sequencing-based omics has transformed our understanding of the gut microbiome^1,2^. But these approaches provide limited information about cellular traits such as size and morphology. Giant bacteria remain overlooked in gut microbiome research, despite their extraordinary cell size, extreme polyploidy and unusual reproductive strategies^3–5^. *Epulopiscium*, the largest known heterotrophic bacteria, has been studied primarily as an intestinal symbiont of surgeonfish, leaving its broader host range unresolved^6–10^. Here we show, through a large-scale analysis of publicly available 16S ribosomal RNA amplicon samples, additional sequencing, and fluorescence microscopy, that *Epulopiscium* spp. occur in humans and many other animals. Phylogenetic analysis resolved multiple *Epulopiscium*-related lineages with distinct host compositions. In humans, *Epulopiscium* sequences were most abundant in infants. In adults, abundance varied across diseases, increasing in inflammatory bowel disease and colorectal cancer while decreasing in neurological conditions compared with healthy adults. Similar patterns were also observed in mouse models. Human *Epulopiscium* cells were 20-70 µm long with morphological traits of other *Epulopiscium* spp. These findings establish giant bacteria as an overlooked component of gut microbiome diversity and highlight how combining sequencing with classical microbiological approaches such as microscopy can uncover organisms with distinctive cellular traits that are difficult to recognise from sequence data alone.

## Main

Over the past two decades, culture-independent sequencing approaches have transformed the study of the human gut microbiome, advancing from broad surveys of community composition to genome analyses and longitudinal studies that integrate multiple omics layers to examine interactions between hosts and microbes^1,11^. Large-scale initiatives have catalogued the taxonomic and functional diversity of gut microbial communities, while increasingly comprehensive reference genome collections enable characterisation at species and strain resolution across thousands of microbial lineages^11^. Nevertheless, the field remains dominated by sequence-based measurements. Standard microbiome workflows generally quantify relative abundances inferred from DNA and are susceptible to biases introduced by experimental protocols, while providing little direct information about cellular traits such as size, morphology and biomass^12–14^. Consequently, unusual phenotypic or physiological traits may remain unrecognised even when the organisms carrying them are detected in microbiome datasets. Giant bacteria represent one potential blind spot in current studies of the human gut microbiome^15^. Their unusually large cell size may make them particularly susceptible to size-selective loss during sample preparation, as most gut microbiome workflows use filtration and centrifugation to remove particulate material before downstream analysis^7,16^.

Giant bacteria challenge the apparent boundary between bacterial organisation and cell size. Bacteria lack dedicated intracellular transport systems and rely on stochastic molecular diffusion to distribute essential proteins and metabolites within the cell^17^. This strategy remains effective over distances of approximately 1 μm, but becomes increasingly inefficient at larger spatial scales^7^. Giant bacteria extend far beyond the conventional bacterial size range, with some attaining centimetre-scale lengths^7,18^. Many giant bacteria are extremely polyploid, a feature thought to distribute genomic templates throughout their large cellular volumes and thereby mitigate diffusion constraints^4,7,19^. Additionally, in non-filamentous giant bacteria like *Achromatium* and *Epulopiscium*, significant allelic divergence across chromosomes has been observed^7,10,20^. Some giant bacteria also possess elaborate intracellular organisation and compartmentalisation, challenging the traditional view of bacterial cells as structurally simple entities^21^. These departures extend to reproduction, and giant bacteria like *Epulopiscium* generate non-dormant offspring within the mother cell through developmental programmes thought to have evolved from endospore formation^5^.

*Epulopiscium*, first described in tropical surgeonfish, is the largest known heterotrophic bacterium^22^. The largest morphotypes can reach more than 600 µm in length^8^. Physiological studies conducted in fish suggested that *Epulopiscium* can alter intestinal pH and host digestive enzyme activity^9^. Genomic and transcriptomic analyses further implicate these bacteria in the degradation and fermentation of complex dietary algal polysaccharides, potentially making otherwise inaccessible nutrients available to their hosts^23,24^. In tropical surgeonfish, more than 90% of the predicted enzymes involved in algal polysaccharide degradation were attributed to *Epulopiscium* and related giant bacteria^23^. Moreover, the association of *Epulopiscium* distribution with host identity and diet, together with the congruence between bacterial and surgeonfish phylogenies, is consistent with long-term ecological specialisation and possible coevolution with their hosts^8,25^.

*Epulopiscium viviparus* produces two metabolically active offspring that inherit only a small fraction of the tens of thousands of chromosomes of the mother cell, while most of the remaining maternal DNA is degraded before maternal cell lysis^3^. Furthermore, to maintain genetic diversity and limit the accumulation of deleterious mutations through Muller’s ratchet, the cells employ horizontal uptake of lysed maternal DNA in the sympatric gut population followed by homologous recombination^10^. *Epulopiscium* displays substantial phylogenetic diversity and has been associated with multiple surgeonfish species, yet its broader host range and geographical distribution remain poorly resolved^8^.

Here we show that bacteria phylogenetically identified as *Epulopiscium* are present in the gut microbiomes of humans and across the animal kingdom. In humans, its abundance (sequence frequency) is associated with known medical conditions. We further propose that common sample preparation protocols, which include filtration or centrifugation to remove large debris from stool samples, bias against giant bacteria, leaving these undetected and systematically underassessed.

## Results

### *Epulopiscium* occurs in the human gut

We collected and analysed 542,340 16S rRNA public amplicon samples from the National Center for Biotechnology Information (NCBI) Sequence Read Archive (SRA). Of these, 247,782 were human, 128,158 were mouse models, and 166,400 were other animals. Given the limited methodological details available in public SRA metadata and the potential bias of sample-processing procedures, such as filtration and centrifugation, towards the preferential recovery of smaller bacterial cells^26–28^, sequence relative abundances were analysed only in *Epulopiscium*-positive (n+) samples. This likely hides true absences, minimizing the significance of the detected enrichments.

The human samples covered 83 countries across all continents, comprising 886 independent studies (Extended Data Fig. 1a). Most studies were from North America, Europe and East Asia, reflecting the current landscape of gut microbiome research (Extended Data Fig. 1b). *Epulopiscium*-positive samples were detected in all defined geographical regions.

*Epulopiscium* was detected across all age groups, but its relative abundance differed markedly among them (Fig. 1b). *Epulopiscium* abundance was significantly higher in both the 0-6 month and 6-12 month groups compared to all other age groups. In contrast, *Epulopiscium* abundance remained broadly comparable among children older than one year, adults, and older individuals, with no significant differences detected.

**Figure 1.**
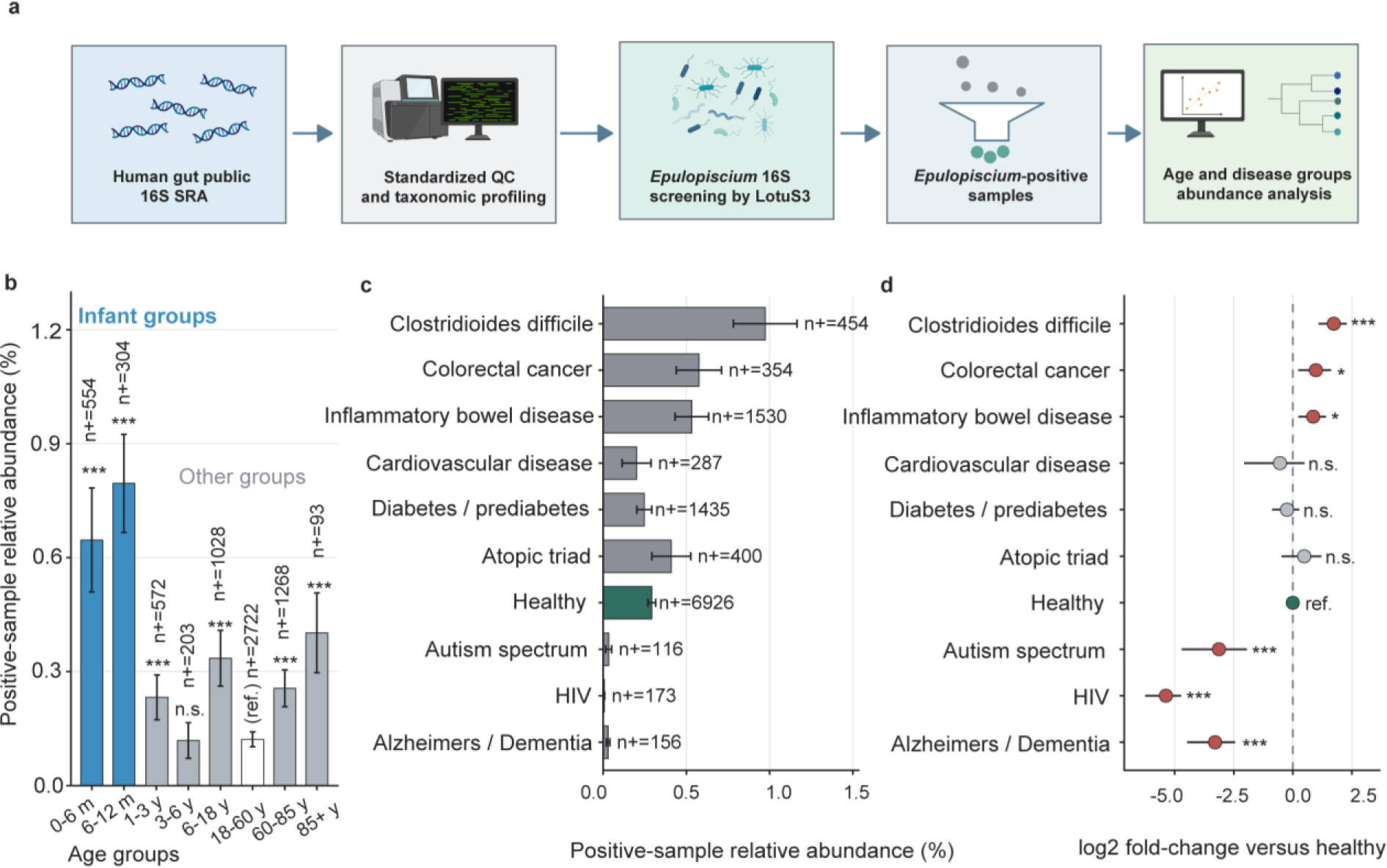
*Epulopiscium* across human age groups and disease-related gut microbiome states. a) Schematic overview of the analysis of public human gut 16S rRNA datasets. Samples underwent standardized quality control and taxonomic profiling, followed by Epulopiscium 16S screening using LotuS3. Epulopiscium-positive samples were subsequently grouped by age and health condition for abundance (sequence frequency) analysis. b) Relative abundance of Epulopiscium among positive samples across age groups. The number of Epulopiscium-positive samples (n+) is indicated for each age group. c) Relative abundance of Epulopiscium among Epulopiscium-positive samples across human health and selected disease conditions. Bars show positive sample relative abundance, with the number of Epulopiscium-positive samples (n+) indicated for each group. d) Log2 fold-change in Epulopiscium relative abundance across health and selected disease conditions compared with healthy controls. Points indicate estimated fold-changes and horizontal lines indicate the corresponding confidence intervals. Statistical significance relative to the healthy reference group is indicated as not significant (n.s.), * (p<0.05), ** (p<0.01) or *** (p<0.001), with healthy shown as the reference.

Across multiple independent studies, *Epulopiscium* showed contrasting distribution patterns between healthy individuals and several disease conditions (Fig. 1c), including intestinal inflammatory and neurological disease settings. These comparisons were based on samples from multiple BioProjects, both healthy and disease-associated. Notably, significant enrichment of *Epulopiscium* was observed in colorectal cancer (CRC), inflammatory bowel disease (IBD) and *Clostridioides difficile* infection (CDI) (Fig. 1d), three intestinal disease contexts in which perturbation of the gut microbiome has been extensively reported^29–31^. In contrast, *Epulopiscium* abundance was significantly lower in individuals with HIV than in healthy controls. Reduced *Epulopiscium* abundance was also observed in autism spectrum disorder and Alzheimer’s disease (Fig. 1d), two neurological disorders that have been consistently associated with alterations in gut microbial composition compared with healthy individuals^32,33^. Within individual BioProjects, comparisons of *Epulopiscium* relative abundance between disease groups and healthy controls were often limited by the absence of detectable *Epulopiscium* in healthy control samples, precluding such comparisons among *Epulopiscium*-positive samples in individual studies.

### *Epulopiscium* has a broad host range

We further asked whether this directional contrast observed in humans was also evident in the mouse model, the most widely used animal model in gut microbiome research. We examined the metadata from 255 BioProjects with mouse models in gut research (Fig. 2a). *Epulopiscium* were similarly enriched in IBD mouse models and depleted in neurological or neuropsychiatric disease models, mirroring the contrasting patterns observed in human datasets. Notably, *Epulopiscium* were significantly more abundant in the guts of genetically engineered and humanised mice than in any other disease model.

**Figure 2.**
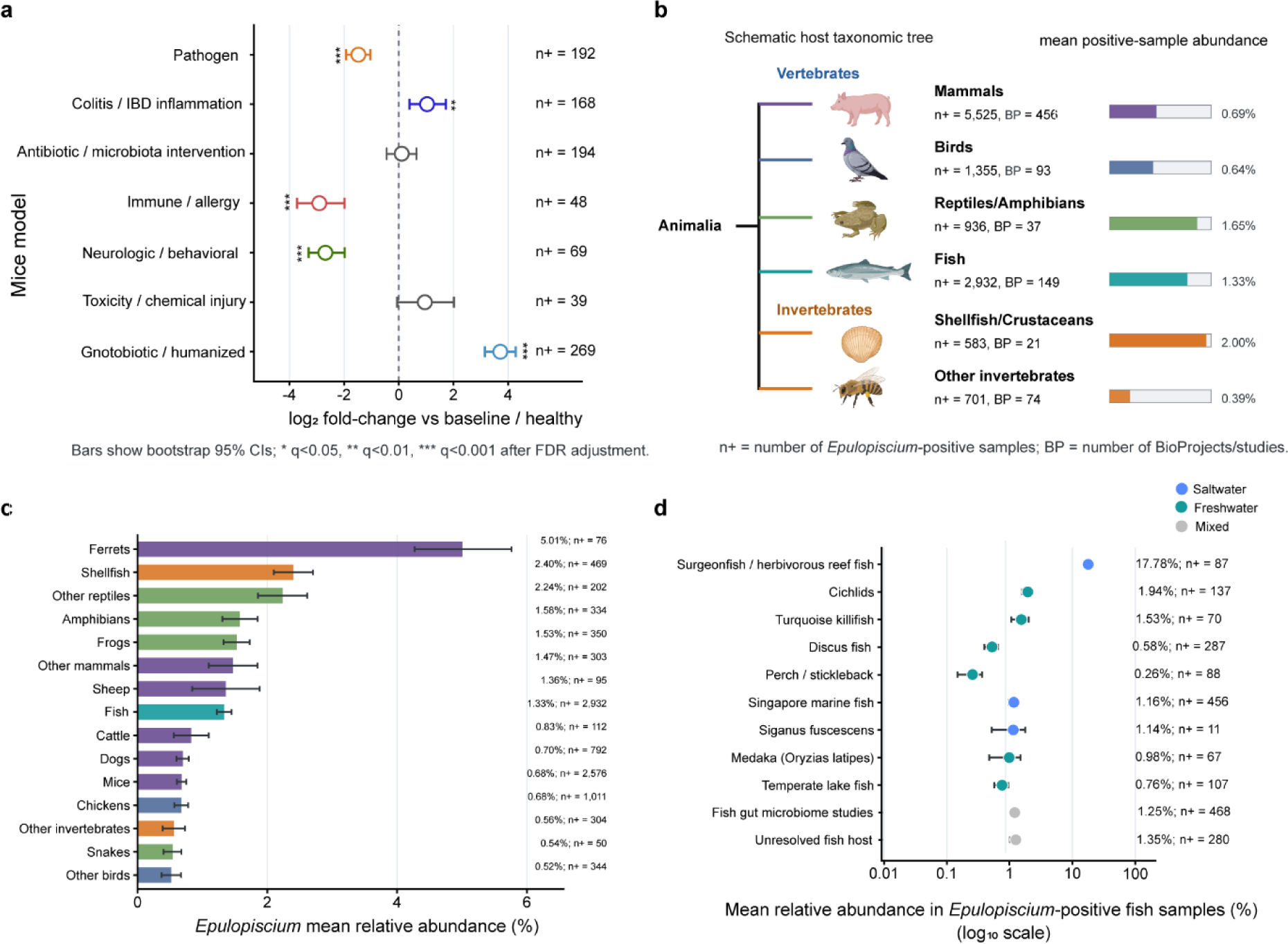
*Epulopiscium* across diverse animal hosts. **a,** Relative abundance of *Epulopiscium* across mouse experimental model categories compared with baseline or healthy controls. Points indicate log2 fold-changes and horizontal lines indicate bootstrap 95% confidence intervals. The number of *Epulopiscium*-positive samples (n+) is shown for each category. Statistical significance after FDR correction is indicated by q values (* < 0.05, ** < 0.01 and *** < 0.001). **b,** Schematic overview of the distribution of *Epulopiscium* across animal groups. Tip size indicates the number of *Epulopiscium*-positive samples (n+), and bars show the mean relative abundance among positive samples. The number of independent BioProjects (BP) is indicated for each host group. **c,** Mean relative abundance of *Epulopiscium* among positive samples across animal host groups. Bars show mean relative abundance, with the number of positive samples (n+) indicated for each group. **d**, Mean relative abundance of *Epulopiscium* among positive fish samples across fish host groups. Points indicate mean relative abundance on a log10 scale, with horizontal error bars showing the corresponding uncertainty and the number of positive samples (n+) indicated for each group.

More broadly, we extended our analysis across a broader range of animal hosts to investigate how widely giant bacteria of the genus *Epulopiscium* are distributed in nature. Across the summarised animal categories, we identified 12,032 *Epulopiscium*-positive samples across 830 BioProjects with retrievable metadata (Fig. 2b). Beyond its canonical fish hosts, we identified *Epulopiscium* across a wide range of animal gut microbiomes, substantially expanding its known host range. The highest mean positive-sample abundances among displayed terminal host groups are ferrets (5.01%, n+=76) and shellfish (2.40%, n+=469) (Fig. 2c).

Because the giant bacterium *Epulopiscium* was originally discovered in surgeonfish, we examined its presence across a broader range of fish species to determine whether this lineage is also present beyond its typical host (Fig. 2d). *Epulopiscium* was detected across diverse fish species, yet its highest relative abundance was observed in its canonical host, surgeonfish (17.78% n+=87). By contrast, several fish categories, including *Sparus aurata* (n+=0/120) and *Oncorhynchus mykiss* (n+=0/65), showed no detectable *Epulopiscium*-positive samples.

### Distinct human *Epulopiscium* lineages

Abundance (sequence frequency) analyses showed that *Epulopiscium* is broadly distributed across human and animal gut microbiomes. However, taxonomic profiles derived from short 16S rRNA gene amplicons could not resolve whether the signals detected in different hosts represented closely related or phylogenetically distinct lineages. Therefore, we analysed publicly available full-length 16S rRNA gene sequences from the studies included in our survey, together with reference sequences from the SILVA ribosomal RNA gene database (v. 138.2). In addition, we generated full-length 16S rRNA gene sequences from patients with systemic lupus erythematosus (SLE) and healthy individuals to resolve the phylogenetic placement and diversity of *Epulopiscium* (Fig. 3).

**Figure 3.**
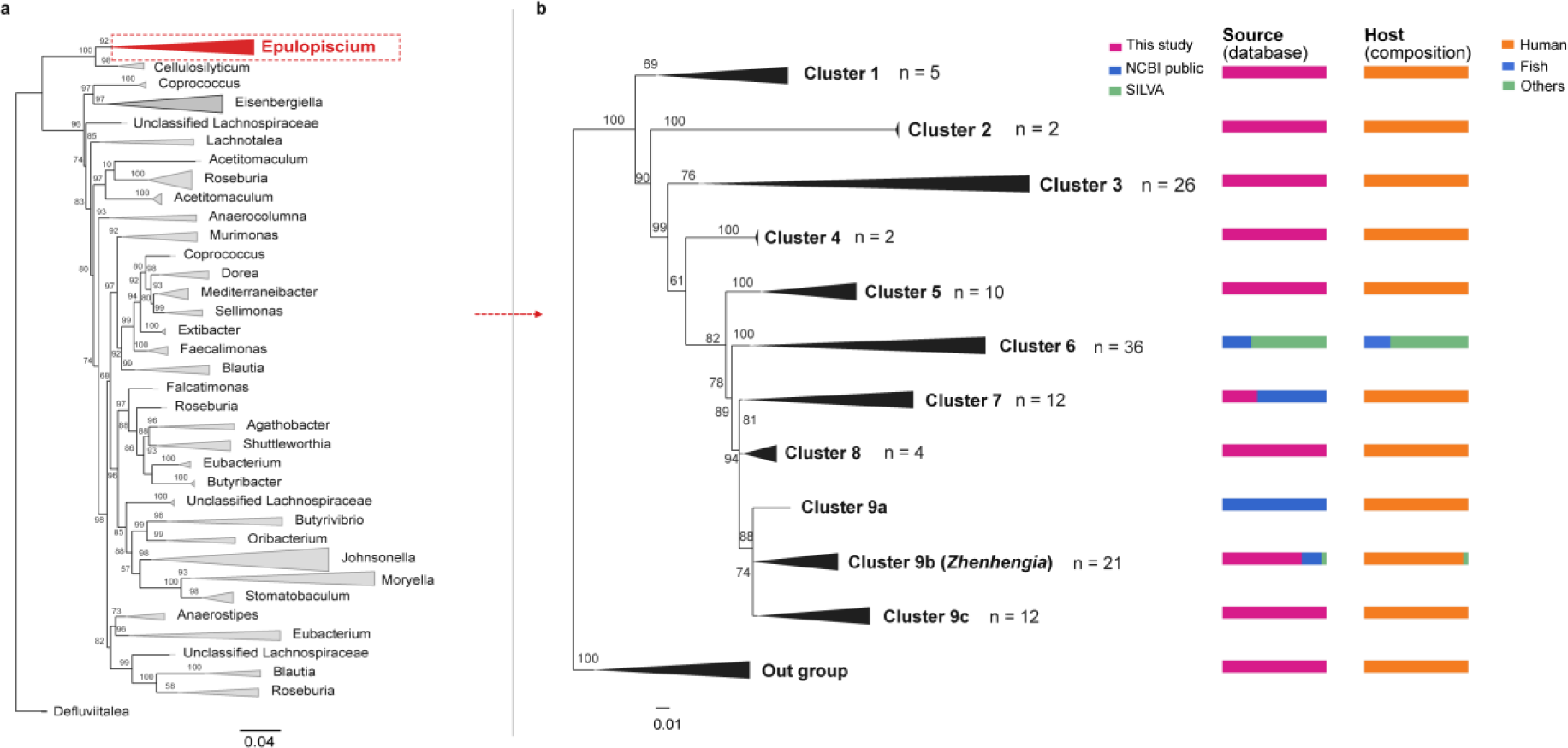
Phylogenetic placement of *Epulopiscium* in the *Lachnospiraceae* family. a, Overview of approximate maximum-likelihood phylogram, comprising 774 focal and *Lachnospiraceae* reference sequences. The tree contains SILVA Epulopiscium references, public OTUs from NCBI and candidate sequences generated in this study. Internal node numbers indicate Shimodaira–Hasegawa-like local support values. b, Rooted maximum-likelihood phylogram inferred with IQ-TREE 2 from 132 selected full-length 16S rRNA sequences using the GTR+F+R5 model and an outgroup for rooting. Collapsed triangles denote visualisation clusters rather than formal taxonomic groups. Cluster 9 was expanded into three subclusters. *Zhenhengia yiwuensis* is highlighted in cluster 9b because its 16S rRNA sequence was assigned to *Epulopiscium* in SILVA classification, although it has not been reported to exhibit giant cell morphology. The outgroup is *Clostridium perfringens.* Internal node values indicate ultrafast bootstrap support (UFBoot: 1,000 replicates) and only UFBoot values >50 are shown. Aligned bars summarise sequence source (this study, public NCBI datasets or SILVA) and host composition (human, fish or others).

The phylogenetic dataset comprised 132 full-length 16S rRNA gene sequences that met the length and quality criteria for phylogenetic analysis, including 83 generated in this study, 23 retrieved from public SRA datasets, 25 SILVA *Epulopiscium* reference sequences, and 1 outgroup. At the family level, the candidates and reference sequences formed a distinct *Epulopiscium*-related clade within the *Lachnospiraceae*, separated from *Cellulosilyticum*, *Coprococcus* and other related genera (Fig. 3a).

Expansion of the *Epulopiscium* clade resolved multiple phylogenetic clusters with distinct source and host compositions (Fig. 3b). Human-derived sequences were distributed across most of these clusters, and sequences generated in this study contributed to several phylogenetically separated lineages. By contrast, cluster 6 comprised public and SILVA sequences derived from fish and other hosts. Deeper phylogenetic relationships received relatively low SH-aLRT and UFBoot support, precluding precise inference of their branching order. Notably, cluster 9b included *Zhenhengia yiwuensis*, a small-cell sized, cultured member of the *Lachnospiraceae* classified under *Epulopiscium* in SILVA taxonomy, which we highlighted as an additional taxonomic reference within this lineage. This indicates that not all members of this lineage necessarily exhibit giant cell morphology. Nevertheless, the full-length phylogeny independently supported the assignment of the detected sequences to *Epulopiscium*-related lineages, thereby providing further evidence for the presence of *Epulopiscium* in the human and animal gut microbiomes.

### Microscopic visualization of *Epulopiscium* in the human gut

To further investigate the presence of *Epulopiscium* and related giant bacteria in the human gut microbiome, we collected clinical stool samples and divided them into two aliquots. One aliquot was used for 16S rRNA gene amplicon sequencing without prior filtration, while the other was retained for microscopy. For samples testing positive for *Epulopiscium*-related sequences, stool from the corresponding aliquot was washed and subsequently filtered through a 10 µm membrane to enrich for large bacterial cells by removing most conventionally sized bacteria.

Bright-field microscopy revealed that the largest observed cell exceeded 65 µm in length and 5 µm in width, with morphological features closely resembling those of *Epulopiscium* previously described from surgeonfish (Fig. 4a)^34^. Other observed cells ranged from 15 to 60 µm in length and from 5 to 10 µm in width (Fig. 4e, i; Extended data Fig. 4). Visible intracellular structures resembling developing offspring cells were observed in Cell 2 (Fig. 4e-h). Notably, some elongated cells exhibited localized protrusions and morphological irregularities near one end (Extended data Fig. 4, cells 5-8). Similar morphological features have previously been documented in *Epulopiscium* sp. type B^35^, where three-dimensional confocal reconstruction revealed a daughter cell emerging through a rupture in the mother cell envelope (Fig. 2L of ref. 35). Giant bacterial cells that were not labeled with the specific *Epulopiscium* probes were also observed (Extended Data Fig. 4, Cell 10). Control experiments using *Escherichia coli* and natural communities of small-sized gut bacteria supported the specificity of the probes (Extended Data Fig. 5). DAPI staining and FISH with the bacterial probe EUB338-I further supported the morphological observations and the Bacterial nature of these cells. FISH with the EPU-mix probes supported their taxonomic identification as *Epulopiscium* (Fig. 4).

**Figure 4.**
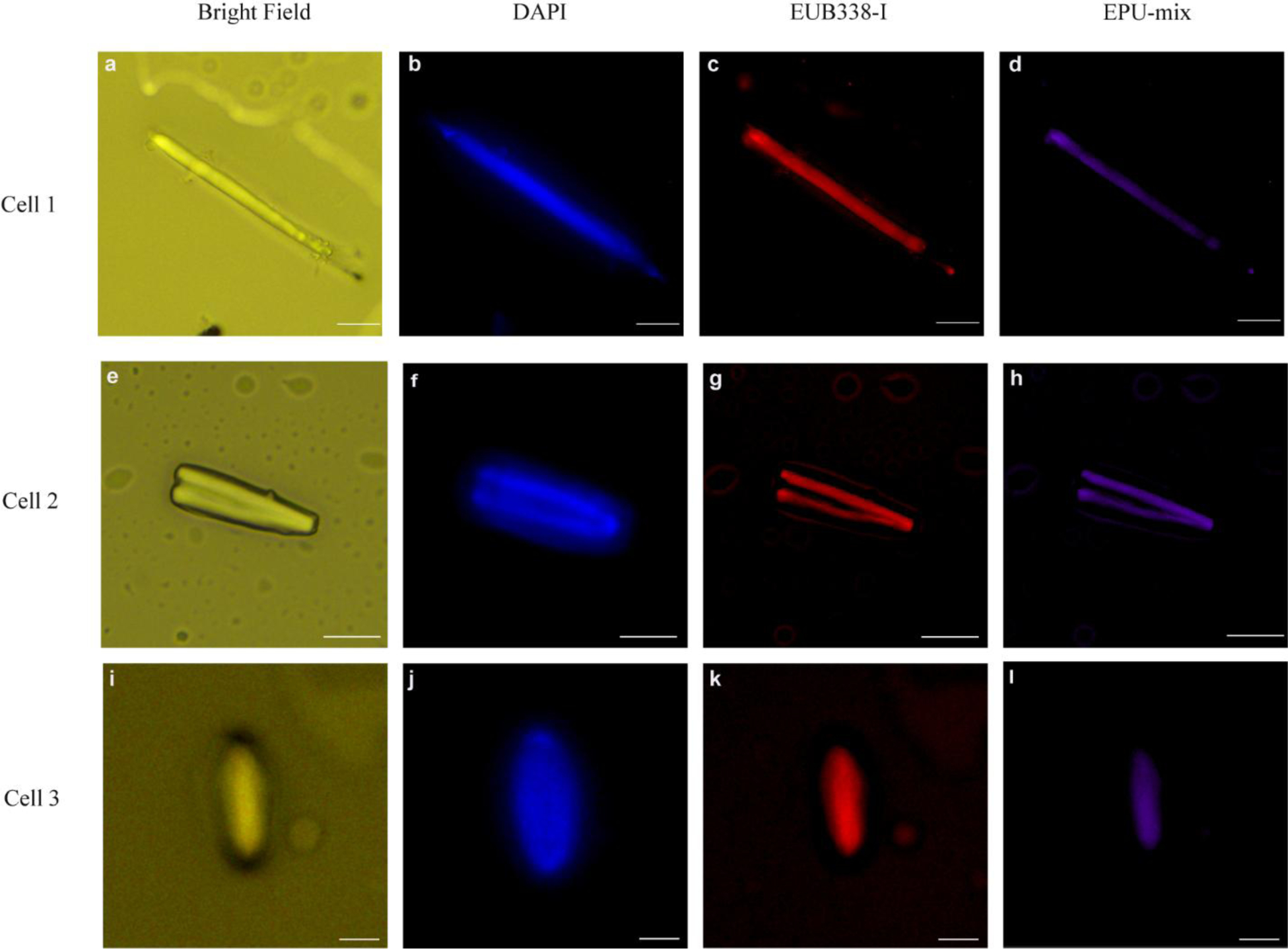
Microscopic images of *Epulopiscium* from humans. Representative microscopy images of three bacterial cells with distinct morphologies. a–d, An elongated cell measuring over 65 μm in length and 5 μm in width; e–h, A cell containing two distinct internal structures suggestive of developing daughter cells; i–l, A smaller, cigar-shaped cell. Bright-field images (a, e, i), DAPI staining (blue; b, f, j), and fluorescence in situ hybridization (FISH) with the bacterial probe EUB338-I (red; c, g, k) and *Epulopiscium*-targeting probes (EPU-mix, magenta; d, h, l) are shown. Scale bars, 10 μm (a–h) and 5 μm (i–l).

## Discussion

Our findings reveal giant bacteria in the human gut, associate their abundance with medical conditions, and show how sample pretreatment can exclude these organisms from microbiome surveys. FISH and microscopy provide direct cellular evidence for giant *Epulopiscium*-related bacteria in human faecal samples, complemented by full-length 16S rRNA gene sequencing and phylogenetic analyses that place human-derived sequences within multiple related lineages. Analysis of more than 500,000 publicly available human (Fig. 1) and other animal samples (Fig. 2) establishes the broader distribution of these bacteria and their associations with human medical conditions. Together, these findings identify an overlooked component of the human gut microbiota and highlight how sample preparation can obscure taxa with atypical cell sizes and morphologies.

Among sequence-positive samples, relative abundance of *Epulopiscium* sequences varied with age and clinical context (Fig. 1). The highest abundances occurred during the first year of life, when gut communities are still assembling and differ from adult communities in nutrient availability, immune maturation and ecological complexity^36,37^. Higher abundances were also observed in CDI, CRC and IBD, whereas lower abundances occurred in HIV, autism-spectrum disorder and Alzheimer’s disease datasets. Directionally similar patterns in several mouse-model categories provide a basis for investigating whether some of these associations reflect shared ecological conditions across hosts. Nevertheless, the heterogeneous responses across clinical settings suggest an association with particular consequences of ecological change, rather than with perturbation in general. Changes in nutrient availability, immune pressure or community structure offer possible explanations^38,39^. Consistent with a broader change in community composition, higher *Epulopiscium* relative abundance was associated with lower relative abundance of a shared set of 16 genera across CRC, IBD and CDI, including *Anaerobutyricum*, *Eubacterium* and *Veillonella* (Extended Data Fig. 3). Studies on fish gut microbiomes have suggested that *Epulopiscium* can influence the activities of intestinal amylases, proteases, and lipases, thereby affecting the host’s capacity to utilize nutrients^9^. In addition, enzymes encoded by *Epulopiscium* contribute to the metabolism of certain nutrients within the host gut^40^. These studies provide valuable insights, but how giant bacteria influence the host and other members of the gut microbiota in the human intestine remains to be further investigated. To distinguish variation in the abundance of detected *Epulopiscium* signals from variation in detection frequency, we restricted abundance comparisons to sequence-positive samples (Figs. 1 and 2). Non-detection in these heterogeneous public datasets could reflect biological absence, limited sequencing sensitivity or reduced recovery during sample processing, potentially including size-selective loss of giant cells. The available metadata did not allow these possibilities to be distinguished reliably.

Our full-length 16S rRNA gene phylogenies further revealed multiple clusters with differing host compositions, with human-derived sequences distributed across most of these clusters rather than confined to a single branch (Fig. 3). Thus, the occurrence of *Epulopiscium*-related bacteria in humans encompasses multiple lineages, raising the possibility that the ecological patterns observed at the genus level reflect contributions from organisms with distinct host associations or ecological requirements. Such analyses would help distinguish ecological features shared across the group from those associated with particular lineages and host environments. Previous studies have shown that, in fish, the distribution of *Epulopiscium* may vary among different host species^8^.

The cellular observations also have implications for interpreting sequence-based abundance. *Epulopiscium* has previously been shown to exhibit extreme polyploidy^41^, suggesting that sequencing-based abundance estimates may not accurately reflect its true biomass. Despite this, multiple large *Epulopiscium* cells could be visualized after filtration of only small quantities of stool, despite their low relative abundance in sequencing data. In conventional gut microbiome studies, filtration, centrifugation, and other sample preprocessing steps may selectively remove larger cells, introducing additional uncertainty during sample preparation^42^. This highlights a potential methodological limitation of current gut microbiome research. Notably, despite the potential loss of large bacterial cells during sample preprocessing, the prevalence of *Epulopiscium* was significantly higher in patients with various diseases than in healthy controls also within individual studies. Integrating microscopy-based cell counts with sequencing data could help more accurately quantify the cellular abundance of these giant bacteria and characterize their cellular properties within the host gut. Beyond its implications for sequence-based abundance estimates, extreme polyploidy may also have important biological consequences. A recently published study predicted that the giant bacterium *Epulopiscium* harbors more antibiotic resistance genes than other giant bacteria, potentially conferring a survival advantage under certain conditions^43^. In *Achromatium*, a large sediment bacterium, extreme polyploidy is accompanied by extensive allelic divergence among chromosome copies, generating substantial genetic diversity within individual cells that has been proposed to facilitate rapid adaptation to changing environmental conditions^20^. The latter has been demonstrated in a recent proteomics study^44^. Although comparable intracellular heterozygosity has not yet been established for *Epulopiscium*, it has been suggested based on existing genomes^7^. Additionally, recombination contributes substantially to population diversification in *Epulopiscium* sp. type B, potentially providing another route for maintaining genetic variation in heterogeneous host environments^10^.

*Epulopiscium* has previously been detected in analyses of human gut microbiome datasets^45–47^ and is also represented in the Human Microbiome Compendium of 168,464 publicly available samples, although its occurrence and biological significance were not mentioned and discussed in the accompanying publication^48^. Sequence-based evidence for its presence in the human gut therefore predates the present study. However, these taxonomic observations have rarely been connected to the organism’s exceptional cellular biology. This gap may partly reflect the scarcity of direct microscopic characterization and the limited integration of environmental and medical microbiological perspectives. Indeed, the identification of *Epulopiscium* in a taxonomic profile does not by itself reveal its giant cell size, extreme polyploidy or unusual reproductive strategy^3,5^.

Our findings provide direct cytological evidence for the presence of giant *Epulopiscium* cells in the human gut and find associations between *Epulopiscium* relative abundance and multiple diseases. Although standardized bioinformatic processing improves comparability across datasets, heterogeneity in study design, sample collection and experimental procedures, together with potential confounding by geography and treatment exposure, still limits comparability.

Taxonomic inventories alone provide an incomplete account of gut microbial diversity. *Epulopiscium* is not the only unusually large bacterium associated with animal digestive systems. *Metabacterium polyspora*, a polysporogenic inhabitant of the guinea pig gut, is closely related to *Epulopiscium*, whereas giant *Spirochaetes, Cristispira* sp. inhabit the digestive structures of molluscs^49,50^, raising the possibility that additional examples remain unrecognized within existing microbiome resources. By combining large-scale sequence surveys with phylogenetic analysis and targeted cellular observation, our study establishes a basis for investigating giant bacteria as a component of human gut-associated microbial diversity. Integrating imaging, quantitative profiling and genome characterization will help determine their abundance and physiological roles. More broadly, incorporating cellular traits alongside taxonomic and genomic information will be important for understanding how unusual microbial forms contribute to host-associated ecology.

## Methods

### Public 16S rRNA data collection and processing

542,340 16S rRNA amplicon sequencing runs were retrieved from the NCBI Sequence Read Archive (SRA) using the NCBI SRA Toolkit (v3.2.1) prefetch and converted to FASTQ format using fasterq-dump with paired-end reads split into separate files where applicable^51,52^.

Raw SSU rRNA gene amplicon sequencing data were processed independently for each SRA accession using LotuS3 (v3.03)^53^. Reads were quality filtered and dereplicated using SDM, clustered de novo into operational taxonomic units (OTUs) at 97% nucleotide identity, followed by read remapping, chimera removal and LULU-based OTU curation^54–56^. Representative OTU sequences were taxonomically annotated against the SILVA 138.2 SSU reference database^57^ using the LotuS3 lowest-common-ancestor procedure. Unclassified OTUs were retained. For abundance analyses, relative abundance was calculated within *Epulopiscium*-positive samples to distinguish the abundance of detected signals from prevalence across all screened samples. Samples containing at least one OTU classified as *Epulopiscium* were defined as *Epulopiscium*-positive. The final figures were assembled using BioRender (BioRender.com), incorporating graphical elements provided by the platform.

### Faecal sample collection

All experiments were approved by the local ethical committee. Faecal samples were collected and immediately stored at −80 °C. The remaining samples used were conserved at −80 °C in the corresponding author’s laboratory.

### DNA extraction and 16S RNA gene sequencing

Total DNA was extracted from faecal samples using the GeneMATRIX Stool DNA Purification Kit (EURx, Gdańsk, Poland) according to the manufacturer’s instructions. In brief, up to 200 mg of faecal material was mixed with lysis buffer and glass beads, incubated at 70°C for 5 min and mechanically disrupted by vigorous vortexing for 10 min. Following centrifugation, PCR inhibitors and cellular debris were removed by PR buffer precipitation. DNA was subsequently purified using spin columns, washed twice and eluted in 50–100 µl of elution buffer. After the extraction, 16S RNA was amplified using barcoded primer pairs with universal adapter sequences and unique individual nucleotide barcodes for multiplex PCR sequencing. The extracted DNA was quantified using a Qubit (Invitrogen), and integrity was checked using agarose gel electrophoresis. Full-length 16S rRNA gene amplicons were used for Oxford Nanopore sequencing. Sequencing libraries were prepared using the Ligation Sequencing Kit V14 (SQK-LSK114; Oxford Nanopore Technologies, Oxford, UK) and sequenced on a PromethION 2 Solo device using an R10.4.1 PromethION flow cell (FLO-PRO114M)^58^. Sequencing was performed for 72 h. Raw signals were basecalled using the high-accuracy 400-bps model implemented in MinKNOW (v25.09.16) with Dorado basecall server (v7.11.2). Reads with a minimum Q score of 9 were retained as passing reads for downstream processing.

### Taxonomic classification of long-read 16S rRNA sequences

Full-length 16S rRNA gene amplicon reads generated by Oxford Nanopore sequencing were taxonomically classified using Kraken2 (v2.17.1) against a Kraken2-formatted database based on the SILVA 138.2 SSU rRNA reference sequence^57,59^.

### Phylogenetic trees

Near-full-length 16S rRNA sequences were aligned using SINA^60^ (v1.7.3) against the SILVA SSU Ref NR99 release 138.2 reference alignment^57^. Alignment compatibility and sequence coverage were inspected, and columns containing >95% gaps were removed before phylogenetic inference. An approximate maximum-likelihood overview tree was inferred using FastTree (v2.2.0) and used for sequence screening and long-branch inspection^61,62^. The resulting branch-length tree was retained for the broad *Lachnospiraceae* overview shown in Fig. 3a; node values in this panel represent FastTree SH-like local support rather than non-parametric bootstrap support. For the focused analysis of *Epulopiscium*-like sequences (Fig. 3b), a subset of 132 sequences was analysed by maximum likelihood using IQ-TREE 2 (v2.4.0) under the GTR+F+R5 substitution model^63^. Branch support was assessed using 1,000 SH-like approximate likelihood-ratio test replicates and 1,000 ultrafast bootstrap replicates (SH-aLRT and UFBoot2, respectively)^61–64^. Only UFBoot percentages are displayed in Fig. 3b. The analysis included sequences generated in this study, public NCBI-derived OTUs, SILVA *Epulopiscium* references, selected *Lachnospiraceae* relatives and external reference sequences. Reference sequences were obtained from the SILVA SSU 138.2 NR99 database and from 16S rRNA genes extracted from publicly available *Epulopiscium* genome assemblies in NCBI.

Trees were inspected and formatted using FigTree (v1.4.5). Nodes were rotated and monophyletic clades were collapsed for visualization. Cluster designations were introduced solely as visual identifiers and do not represent formally defined taxonomic groups.

### Probe design for FISH

To visualize giant bacterial cells in faecal samples, we designed *Epulopiscium*-specific fluorescence in situ hybridization (FISH) probes based on full-length 16S rRNA gene sequences obtained through Oxford Nanopore sequencing and closely related reference sequences retrieved from the SILVA SSU 138.2 database. The sequences were imported into ARB, and oligonucleotide probes ranging from 19 to 22 nucleotides in length were designed using the ARB Probe Design tool. Three probes were selected: Probe EPU718 (5′– GCTACTGGTGTTCCTCCAAATC–3′), EPU175 (5′–TTATGCGGTATTAGCGTTTGT–3′) and EPU888 (5′–GTCGTACTTCCCAGGTGGAATA–3′). All of them were labelled with Cyanine 5 (Cy5). The probes were named using the first three letters of *Epulopiscium* and the starting position of their binding site on the 16S rRNA. The predicted specificity and taxonomic coverage of the designed probes were evaluated in silico using the SILVA Probe Match and Evaluation Tool (TestProbe 3.0) against the SILVA SSU Ref NR99 138.2 database^57^. Probe matches were examined across different taxonomic groups to assess target coverage and identify potential non-target matches. In silico analyses indicated high predicted specificity and coverage of the majority of *Epulopiscium* sequences identified in our dataset. To experimentally assess potential non-specific hybridization, control experiments were performed using *Escherichia coli* as a non-target organism. Fixed *Escherichia coli* (*E. coli*) cells were hybridized with the universal bacterial probe EUB338-I (5′–GCTGCCTCCCGTAGGAGT–3′) labelled with ATTO 594 and the Epu-mix probe (EPU718, EPU175 and EPU888) labelled with Cy5 under (20%, 30%, and 40% formamide) at 46 °C for 2 h (Extended Data Fig. 6).

### Fluorescence in situ hybridization

Fluorescence in situ hybridization was performed as previously described^65^. Briefly, approximately 0.2 g of faecal material was suspended in 15 mL of PBS. 4 mL samples were fixed in 4% paraformaldehyde for 1 h at room temperature, collected on Millipore 10 µm polycarbonate (PC) filters and embedded in 0.2% low-melting-point agarose. Cells were permeabilized with lysozyme at 37 °C for 30 min. Hybridization was performed using the bacterial probe EUB338-I labelled with ATTO 594 and our designed *Epulopiscium* probes labelled with Cy5 at 46 °C for 2 h using 40% formamide. All oligonucleotide probes were synthesized by biomers.net GmbH (Ulm, Germany). Following washing, samples were mounted in CitiFluor AF1 Plus DAPI, with coverslips applied after 10 min. Bright-field and fluorescence images of the same fields of view were acquired using a Leica LMD6 and processed using LAS X (version 3.10.2.30243; Leica Microsystems). The images were pseudocoloured and processed using Fiji (ImageJ)^66^ for visualization.

### Statistical analysis

Unless otherwise indicated, data are presented as the mean ± s.e.m. Statistical analyses were performed using R (v.4.4.3). Relative abundances in *Epulopiscium*-positive samples from the early-life age groups (0–12 months) and the remaining age groups were compared using a two-sided Wilcoxon rank-sum test. Disease-associated effect sizes relative to the healthy reference group were expressed as log2 fold changes, with 95% percentile confidence intervals and empirical P values estimated from 5,000 bootstrap resamples. For mouse-model comparisons, relative abundances were log2-transformed and each category was compared with the baseline or healthy group using a two-sided Welch’s t-test; 95% confidence intervals were estimated from 5,000 bootstrap resamples. P values from multiple comparisons were adjusted using the Benjamini–Hochberg procedure. An FDR-adjusted q < 0.05 was considered statistically significant.

## Data availability

All sequencing data generated in this study have been deposited in the NCBI Sequence Read Archive (SRA) and are publicly available under BioProject accession number PRJNA1537502. Data used for bioinformatic analyses throughout the Article are available for download from Zenodo (10.5281/zenodo.22273977).

## Code availability

All sequencing and analyses were performed using standard, publicly available software packages and established analysis pipelines, as described in the Methods. Any additional information is available from the corresponding authors on reasonable request.

## Acknowledgements

We thank all members of Umbio for the scientific discussions. We also thank the members of the DRFZ (German Rheumatology Research Centre, Berlin, Germany) for their technical assistance with samples. We thank the DRK clinic in Gransee, Brandenburg, Germany, for participating in the study. We thank all the patients who participated in the study.

## Author information

**Authors and Affiliations**

**Department of Environmental Microbiomics, Technische Universität Berlin, Berlin, Germany**

Yiang Lyu, Marit Grüner, Zhen Han, Aron Klein, Mina Bizic & Danny Ionescu

**Core Facility Bioinformatics, BOKU University, Vienna, Austria**

Luca Zoccarato

**Institute of Computational Biology, BOKU University, Vienna, Austria**

Luca Zoccarato

**Faculty of Marine Sciences, Ruppin Academic Centre, Israel**

Alon Daya

**Institute of Biotechnology, Technische Universität Berlin, Berlin, Germany**

Zhen Han, Hyun-Dong Chang

**German Rheumatology Research Centre, Berlin, A Leibniz Institute, Berlin, Germany**

Hyun-Dong Chang

## Author contributions

Y.L. performed the experiments, conducted the bioinformatic analyses and wrote the original draft of the manuscript. M.G., Z.H., A.K., L.Z. and A.D. contributed to the bioinformatic analyses and revised the manuscript. H.-D.C. provided the patient samples and revised the manuscript. M.B. and D.I. supervised the study and provided guidance throughout the project. All of the authors approved the manuscript before submission.

## Extended data figures and tables

**Extended Data Fig. 1.**
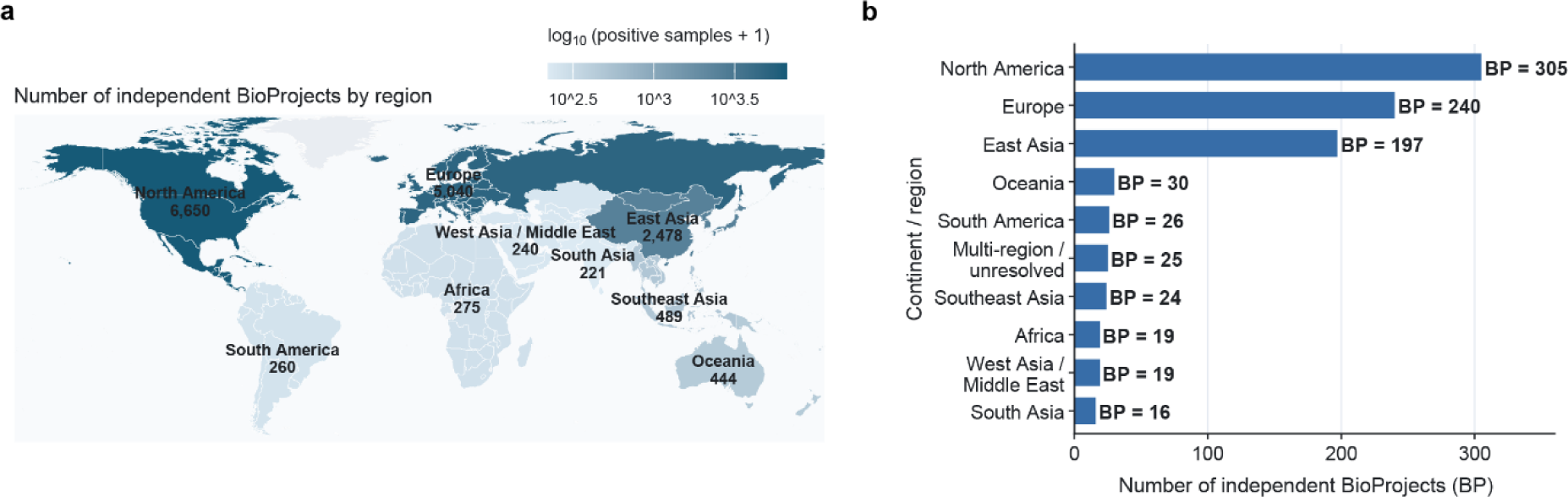
Geographical distribution of human gut microbiome BioProjects and *Epulopiscium*-positive samples. a. World map showing the number of human gut 16S rRNA amplicon samples in which *Epulopiscium* sequences were detected. Labels give the number of positive samples in each geographical region, and the colour scale represents log_10_(n + 1), where n is the number of positive samples. b. Number of BioProjects (BP) by geographical region.

**Extended Data Fig. 2.**
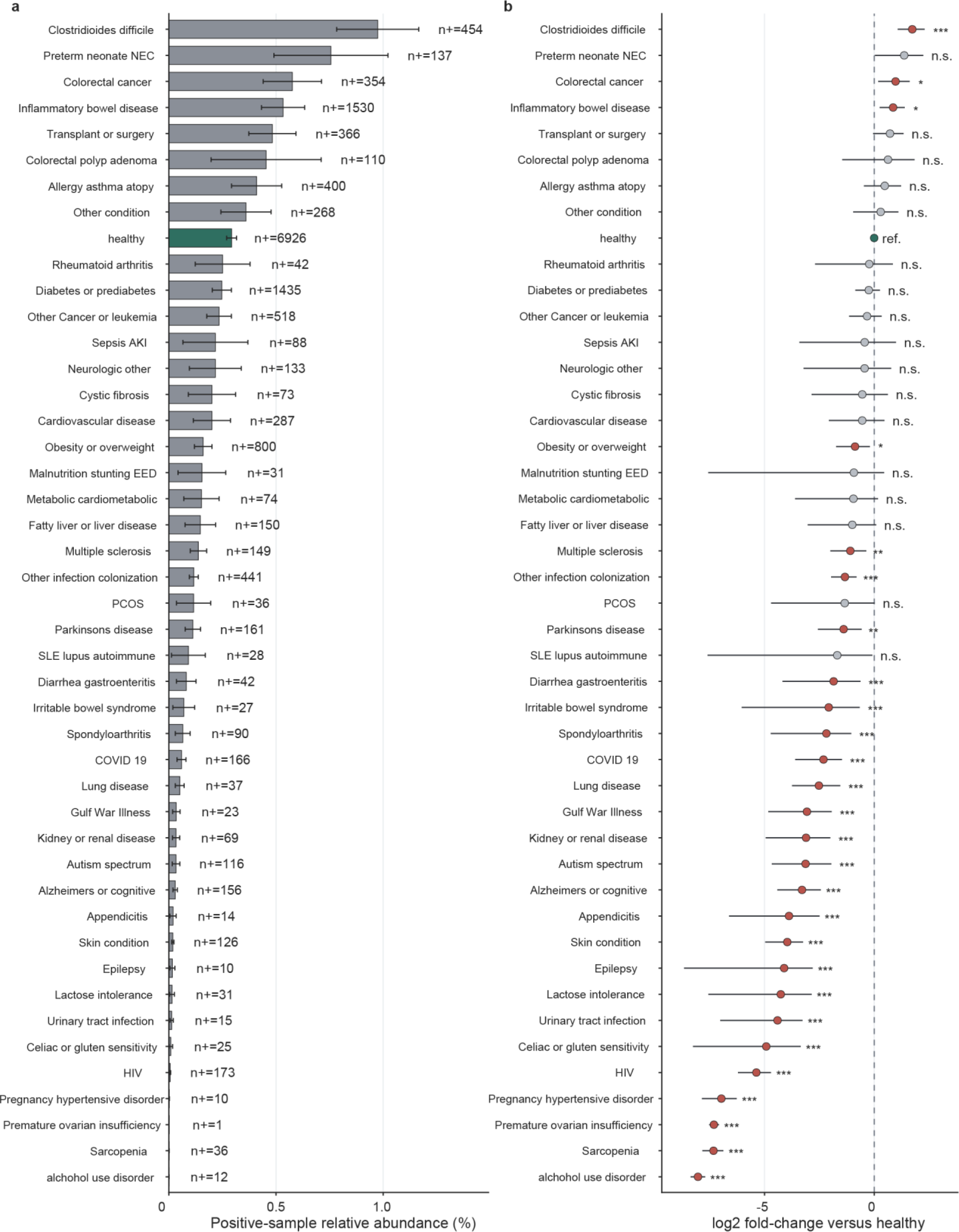
Relative abundance of *Epulopiscium* across human health conditions. a, Mean relative abundance of Epulopiscium in healthy samples and across disease and other clinical categories. Only samples in which these sequences were detected were included. Labels indicate the number of positive samples (n_+_) in each category. b, Log₂ fold changes in relative abundance for each category compared with the healthy reference group (ref.). Points indicate estimated fold changes and horizontal lines show 95% confidence intervals. Confidence intervals and empirical P values were estimated from 5,000 bootstrap resamples. P values were adjusted for multiple comparisons using the Benjamini–Hochberg procedure. n.s., q ≥ 0.05. *q < 0.05, **q < 0.01 and ***q < 0.001.

**Extended Data Fig. 3.**
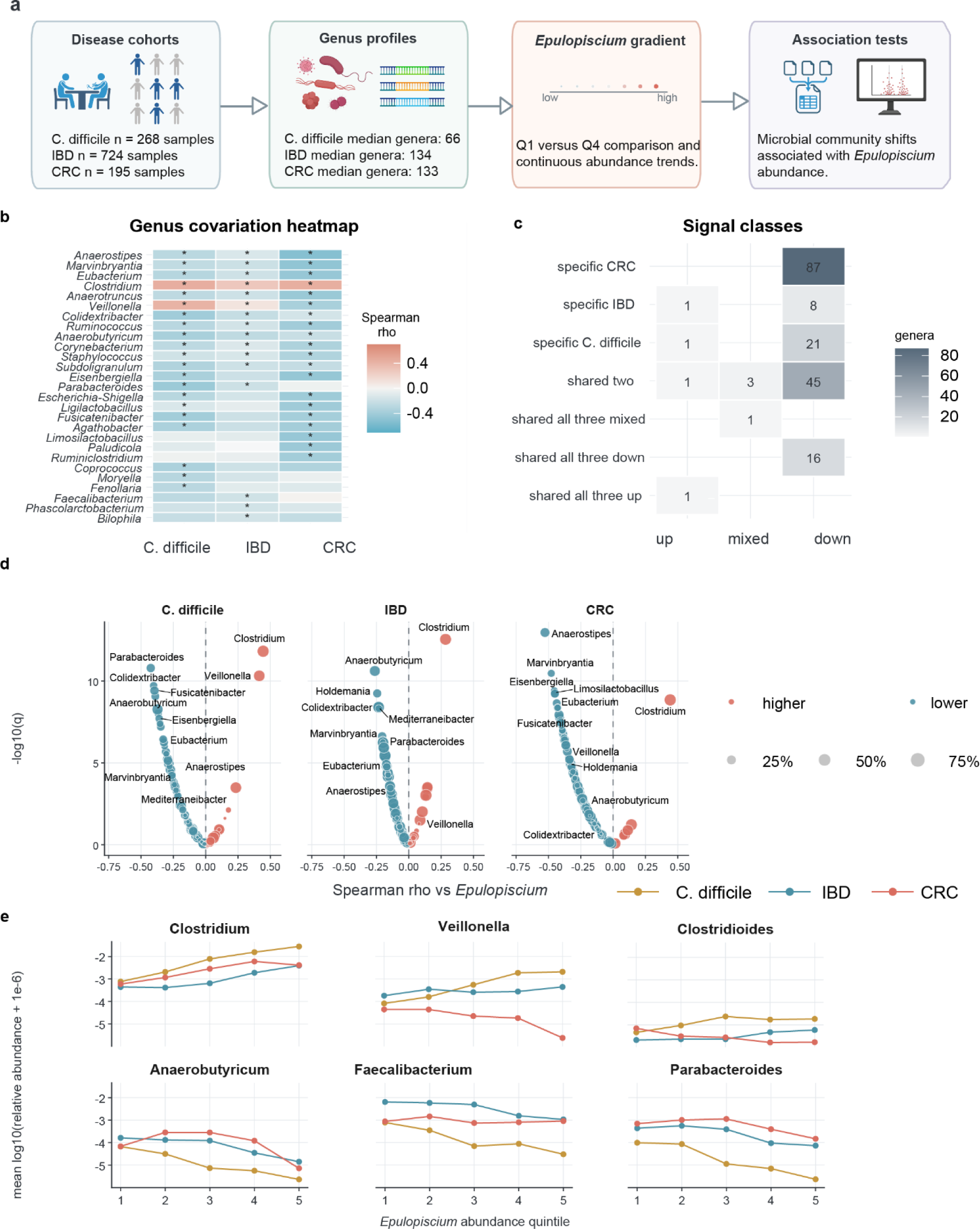
Gut bacterial genera associated with *Epulopiscium*-related sequence abundance across three disease cohorts. a, Overview of genus-level profiling in samples from cohorts with *Clostridioides difficile* infection (CDI, n = 268), inflammatory bowel disease (IBD, n = 724) or colorectal cancer (CRC, n = 195). Analyses compared samples in the lowest and highest quartiles of *Epulopiscium* abundance and assessed associations across the full abundance range. b, Heat map of Spearman’s rank correlation coefficients (ρ) between the relative abundance of selected genera and that of *Epulopiscium* in each cohort. Asterisks indicate significant associations. c, Number of genera with cohort-specific or shared associations, grouped by direction. Sixteen genera showed negative associations in all three cohorts. d, Cohort-specific plots of Spearman’s ρ against −log_10_(q), where q is the multiple-testing-adjusted P value. Each point represents a genus. e, Mean log_10_-transformed relative abundance, calculated as log₁₀(relative abundance + 10^-6^) before averaging, for six selected genera across quintiles of *Epulopiscium* abundance in each cohort.

**Extended Data Fig. 4.**
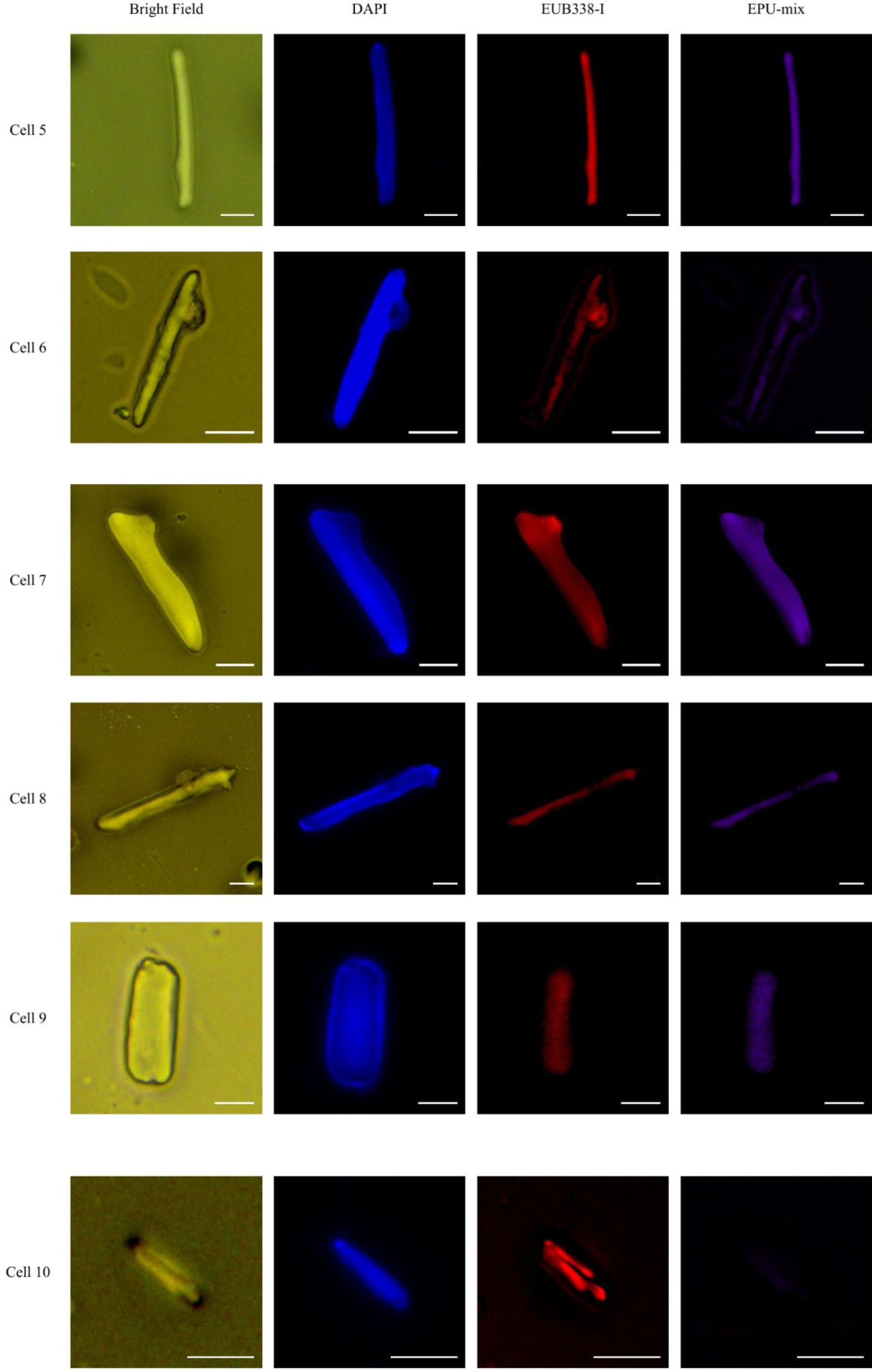
Additional giant bacterial cells in human faecal samples. Representative images of six giant bacterial cells (cells 5–10) shown in bright field and in the DAPI, EUB338-I and EPU-mix fluorescence channels. Scale bars, cells 5–8: 10 µm. Cells 9, 10: 5 µm.

**Extended Data Fig. 5.**
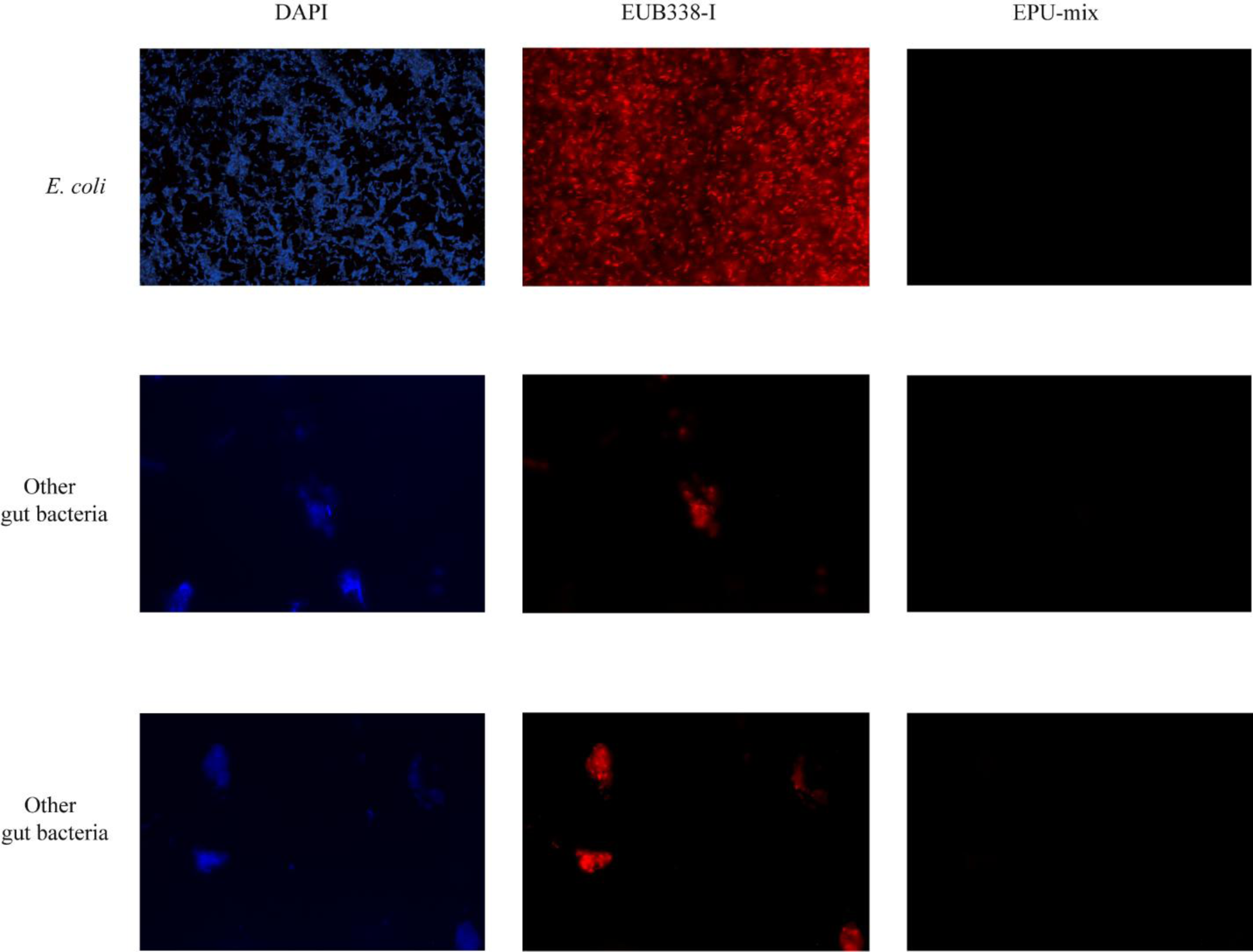
Control images for FISH. Representative images of *Escherichia coli* and other gut bacteria shown in bright field and in the DAPI, EUB338-I and EPU-mix fluorescence channels.

**Extended Data Fig. 6.**
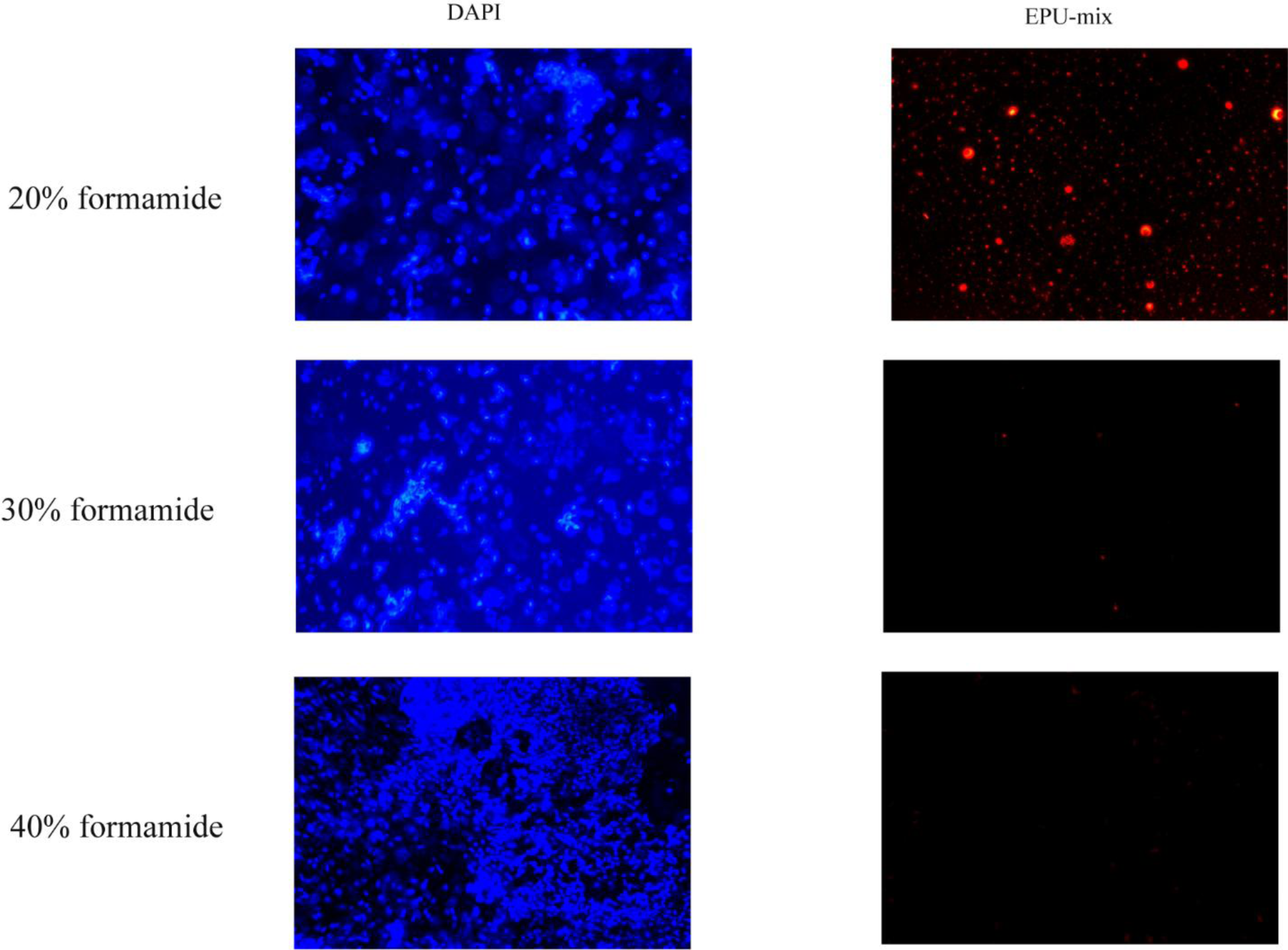
Fluorescence in situ hybridization of *Escherichia coli* with the specific probe set EPU-mix at different formamide concentrations. From top to bottom, the rows show hybridization at 20%, 30%, and 40% formamide. The left column shows DAPI staining (blue), and the right column shows EPU-mix probe fluorescence (red).

